# Kinesin-6 KIF20B is required for efficient cytokinetic furrowing and timely abscission in human cells

**DOI:** 10.1101/171900

**Authors:** Kerstin M. Janisch, Katrina C. McNeely, Joseph M. Dardick, Samuel H. Lim, Noelle D. Dwyer

## Abstract

Cytokinesis requires the cooperation of many cytoskeletal and membrane regulators. Most of the major players required for cytokinesis are known, but the temporal regulation and adaptations for different cell types are less understood. KIF20B (previously called MPHOSPH1 or MPP1) is a member of the Kinesin-6 family, which also includes the better-known members KIF23/MKLP1 and KIF20A/MKLP2. Previously, we showed that mouse Kif20b is involved in cerebral cortex growth and midbody organization of neural stem cells. Here we show that KIF20B has a cell-autonomous role in cytokinesis in isolated human cells. It localizes to microtubules of the central spindle and midbody throughout cytokinesis, at sites distinct from the other Kinesin-6 family members. KIF20B is not required for central spindle or midbody assembly, but affects midbody shape and late maturation steps. KIF20B appears to temporally regulate both furrow ingression and abscission.

## INTRODUCTION

Cytokinesis is fundamentally important to building and renewing tissues, but it is still poorly understood. It consists of two sequential processes: cleavage furrow ingression, which takes minutes, and abscission, which can last more than an hour (for review see Green *et al*., 2012 and Mierzwa and Gerlich, 2014). Moreover, cytokinesis has different spatial and temporal regulation in different cell types or at different times in development (Singh and Pohl, 2014; Lenhart and DiNardo, 2015). While many of the major players of cytokinesis have been defined, how the temporal control of furrowing and abscission is regulated is not well understood. Furthermore, how abscission regulation influences tissue development is only beginning to be explored (Dionne *et al*., 2015).

After chromosome segregation, interpolar microtubules are bundled in an antiparallel manner to form the central spindle, which is involved in specifying and focusing the plane of cleavage. As the cleavage furrow ingresses to form a thin intercellular bridge, the central spindle microtubules are compacted into a dense structure called the midbody. The midbody serves as a platform to mediate the final severing event, abscission. It contains over 150 proteins and lipids (Skop *et al*., 2004; Atilla-Gokcumen *et al*., 2014). During midbody assembly, these proteins are partitioned into distinct subdomains (Elia *et al*., 2011; Green *et al*., 2012; Hu *et al*., 2012). The midbody core or “dark zone” is at the center where microtubules of the two spindle halves overlap in an antiparallel arrangement and are surrounded by electron-dense material (Mullins and Biesele, 1977). PRC1 organizes the antiparallel microtubules in the central spindle and then localizes to the midbody core (Mollinari *et al*., 2002; Hu *et al*., 2012) Around the midbody core is a bulge that appears as a ring by light microscopy. Tightly packed parallel microtubules emanate from either side of the midbody core and form the midbody flanks. The flanks are labelled by AuroraB kinase signal that terminates near the eventual abscission site(s). As the time for abscission gets nearer, constriction sites (also called secondary ingressions) form on one or both sides of the midbody bulge. There the microtubules are even more tightly packed (Mullins and Biesele, 1977). How the constriction sites are positioned and formed is not understood, but ESCRTs and endosomes are involved (Elia *et al*., 2011; Guizetti *et al*., 2011; Schiel *et al*., 2012). After midbody assembly, CEP55 is recruited to the midbody bulge where it scaffolds sequential recruitment of ESCRT-I and –III components on either side of the midbody bulge (Morita *et al*., 2007; Lee *et al*., 2008). Together ESCRT-III filaments and the microtubule-depolymerizing enzyme spastin are thought to complete abscission (Morita *et al*., 2007; Elia *et al*., 2011; Guizetti *et al*., 2011). In HeLa cells and MDCK cells, a second abscission occurs on the other flank to release the midbody (Elia *et al*., 2011; Guizetti *et al*., 2011). Other reports have observed abscission on only one side, and this may depend on cell type or daughter fates (Ettinger *et al*., 2011; Kuo *et al*., 2011).

Microtubule motors are crucial for mediating the cytoskeletal reorganizations that take place during cell division. The Kinesin-6 family members are thought to have roles in cytokinesis and cancer (Baron and Barr, 2015). The family is defined by homology in the motor domain; the stalks and tails are divergent (Dagenbach and Endow, 2004; Miki *et al*., 2005). They are distinguished by a long insertion in loop-6 of the motor domain, and a relatively long neck domain that may enable the two heads in a homodimer to bridge longer distances than adjacent tubulin binding sites. They homodimerize through their coiled-coil domains in the stalks. *C*. *elegans* has one member of this gene family, *zen*-*4*. *Drosophila* has two members, *pavarotti* and *subito*. Vertebrates have three members of the Kinesin-6 family: KIF23/MKLP1, KIF20A/MKLP2, and KIF20B (previously called Mphosph1 or Mpp1). KIF23 partners with mgcRacGAP to promote central spindle assembly (Glotzer, 2005; Nishimura and Yonemura, 2006). Later, KIF23 localizes to the midbody bulge and is required for a stable midbody to form (Matuliene and Kuriyama, 2002; Zhu *et al*., 2005a; Zhu *et al*., 2005b). KIF20A/MKLP2 localizes to the midbody flanks, and is required to recruit AuroraB kinase there (Neef *et al*., 2003; Gruneberg *et al*., 2004; Zhu *et al*., 2005a). KIF20B apparently evolved in the vertebrate lineage to become structurally divergent, with an extraordinarily long stalk (~1000 amino acids) containing 3 hinges that may allow increased flexibility (Kamimoto *et al*., 2001; Dagenbach and Endow, 2004; Miki *et al*., 2005). In cell-free assays, KIF20B acted as a slow, plus-end directed microtubule motor, and was sufficient to slide and bundle microtubules (Abaza *et al*., 2003). When knocked down in human HCT116 cells, a subset of cells failed cytokinesis, and sometimes underwent apoptosis (Abaza *et al*., 2003). However, the role of KIF20B in cytokinesis is not known.

We previously isolated a *Kif20b* mouse mutant in a genetic screen for genes involved in cerebral cortex development (Dwyer *et al*., 2011; Janisch *et al*., 2013). The mutant embryos have small brains (microcephaly) with increased apoptosis. Mutant neurons have wider, more branched axons (McNeely *et al*., 2017). In the embryonic neural stem cells in mutant brains, no changes in mitotic indices or cleavage angles were evident, but abnormalities in cytokinetic midbodies were significant. Midbodies had altered shape and organization at the apical membrane (Janisch *et al*., 2013; Janisch and Dwyer, 2016). We hypothesized that defective or failed abscission in a subset of neural stem cells caused them to undergo apoptosis, thus depleting the progenitor pool, reducing neurogenesis, and leading to a small brain.

Motivated by a desire to understand mechanisms of abscission in polarized neural stem cells, but hindered by the dearth of data about KIF20B’s role in cytokinesis, we set out to investigate it in a simpler more tractable system, to generate hypotheses which can then be tested in primary mouse neural cells and intact brain tissue. In the mutant brains, the observed phenotypes could be due to non cell-autonomous roles of KIF20B, or secondary effects from previous defective divisions. We chose the HeLa human cell line because the cells proliferate as isolated cells, and have a flat morphology, enabling easier imaging of cytokinesis and of the cytoskeleton. They are readily transfected, tolerant of live imaging, and many more antibodies are made against human proteins than mouse. Finally, since the majority of studies of abscission proteins have used HeLa cells, it is an essential baseline cell type for comparison.

We found that KIF20B localizes to microtubules of the central spindle and midbody throughout cytokinesis, in particular near the midbody core and constriction sites, in patterns distinct from the other two Kinesin-6 family members. Depletion of KIF20B in HeLa cells caused some cytokinesis failure as evidenced by increased multinucleate cells. In single cell live imaging, furrow ingression appeared qualitatively normal, but was slower. KIF20B appears not to be required for midbody assembly, but may be involved in regulating midbody width and microtubule constriction sites. Abscission timing was dysregulated and delayed. KIF20B may act in part by stabilizing microtubules. In the context of our previous work, these data suggest that subtle defects in the midbody and temporal control of cytokinesis may have devastating consequences for brain development.

## RESULTS

### KIF20B protein localizes to microtubules in the central spindle and midbody throughout cytokinesis

Before analyzing the specific role of KIF20B in cytokinesis in HeLa cells, we first sought to determine its detailed subcellular localization in relation to the microtubule cytoskeleton during cytokinesis. To do this we coimmunostained HeLa cells for endogenous KIF20B along with alpha-tubulin. KIF20B is a low abundance protein, but it is readily detected in dividing cells when concentrated in cytokinetic structures. In cells fixed in anaphase, KIF20B signal appears as puncta on the microtubules of the central spindle (**Figure 1 A**). In cells with more deeply ingressed furrows, KIF20B is more concentrated in the center of the central spindle as a band (**Figure 1 B**). In early midbodies, KIF20B signal is detected in two discs on either side of the midbody core or “dark zone” (arrowhead) (**Figure 1, C, D**). In late midbodies, KIF20B signal is more extended along the midbody flanks (**Figure 1 E)**. If microtubule constriction sites are visible, KIF20B surrounds them, in this case appearing as four distinct spots on the midbody (**Figure 1 F**). In cells that appear to have undergone abscission (with a large gap in tubulin staining on one side of the midbody), KIF20B can be detected still surrounding the dark zone (arrowheads), and on both flanks on either side of the abscission site (**Figure 1 G – I**). In a few cases where a microtubule strand remained connecting the sister cells, they were decorated with dots of KIF20B (arrows in **Figure 1 G-G”**). Overexpression of a GFP-tagged full-length KIF20B in cell lines or neurons usually causes cell death, as previously reported by our group and others (Abaza *et al*., 2003; McNeely *et al*., 2017). However, a small number of surviving cells expressing GFP-KIF20B showed a similar localization as endogenous KIF20B detected by immunostaining, flanking the dark zone (**Figure 1 J – J”**). Together these data suggest that KIF20B first associates with central spindle microtubules starting in early anaphase, then accumulates at microtubule plus ends that coincide with the plane of furrow ingression, and finally remains concentrated at the narrowest portions of the midbody surrounding the dark zone and constriction sites throughout the abscission process.

**Figure 1.**
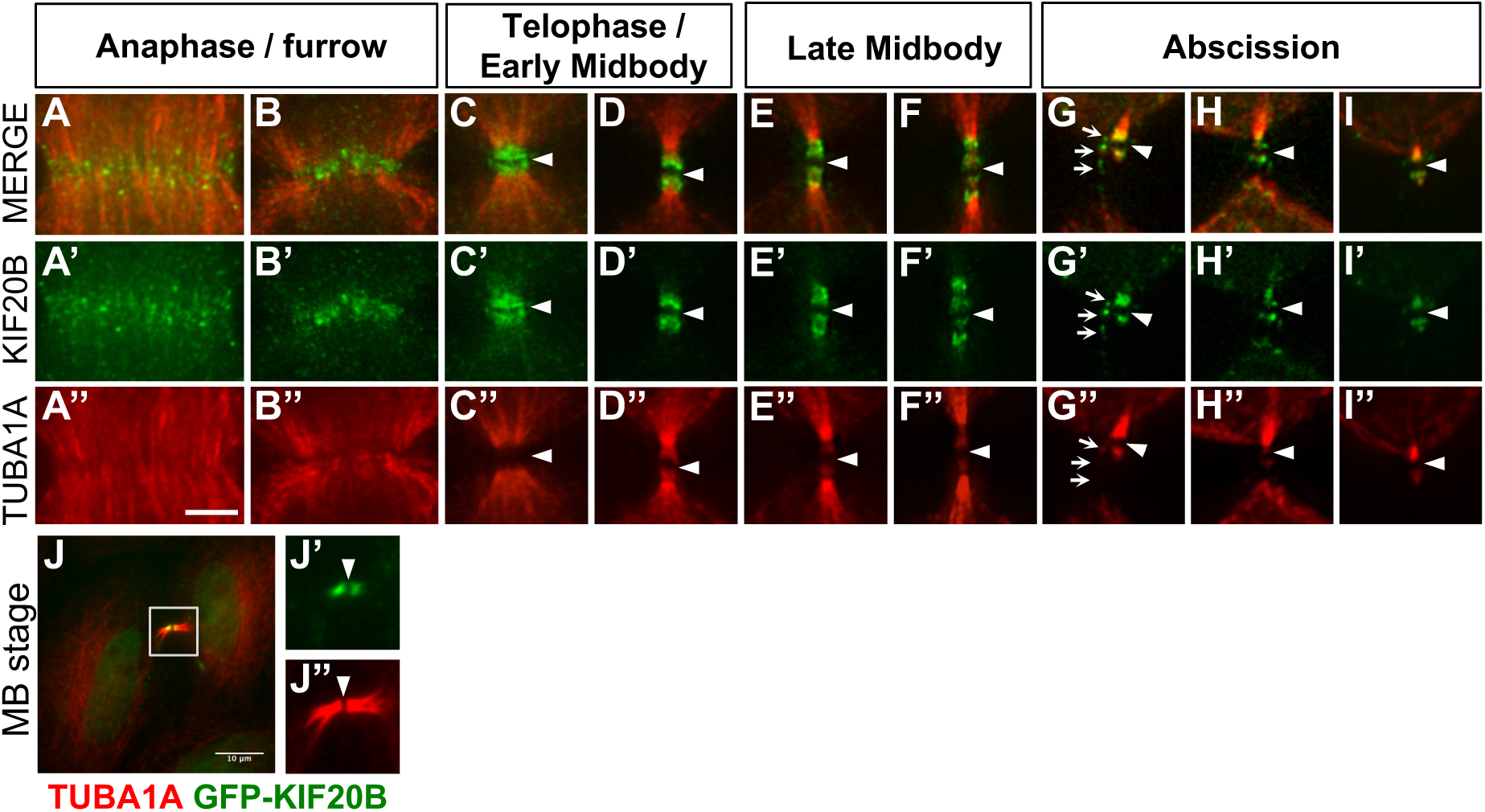
KIF20B localizes to the central spindle and midbody throughout cytokinesis in HeLa cells. (A - B”) During Anaphase, KIF20B starts to accumulate as speckles along the microtubules of the central spindle (A’) forming a dense band in the middle of the central spindle (B’). In Telophase (C - D”), KIF20B accumulates on the inner flanks of the midbody surrounding the dark zone (arrowhead), forming a cap-like structure. In late midbody stage (E - F”), KIF20B spreads out on the midbody flanks surrounding the constriction sites for abscission, resulting in 4 distinct spots of KIF20B localization. After abscission (G - I”), KIF20B can be seen in the remaining midbody connected to one of the daughter cells (G - I’), still surrounding the former dark zone (arrowhead). The small arrows in the G panels point to KIF20B dots localizing along a strand of microtubules. (J - J”) GFP-KIF20B expressed in HeLa cell shows the same localization within the midbody as detected by antibodies to KIF20B. Arrowheads point to dark zone center of the midbody in all pictures. Scale bars are 5 *μ*m for A through I” and 10 *μ*m for panel J.

### Knockdown of KIF20B increases multinucleate cells

To investigate the cell-autonomous requirement and primary role(s) of KIF20B in cell division, we depleted endogenous KIF20B from HeLa cells using small interfering RNA (siRNA) transfections. We used a previously published KIF20B-specific small interference RNA (siRNA) (Abaza *et al*., 2003)and confirmed that it depletes endogenous KIF20B in HeLa cells to undetectable levels by 24 hours post-transfection (**Supplemental Figure 1**). First, we compared general mitotic and cytokinesis parameters between asynchronous cells transfected with control siRNA (siLUC) or siKIF20B (siKIF), and fixed 24 or 48 hours after transfection. Depletion of KIF20B did not significantly change the mitotic index or the midbody index, but caused a slight reduction in telophase cells at 48 hours (**Figure 2 A, B, C**). However, KIF20B depletion resulted in a 2.5-fold increase in the occurrence of multinucleated cells (with two or more clearly distinct nuclei) at 24 hours (**Figure 2 D, 2 F and F’**), and a striking increase in multi-lobed nuclei at 48 hours post-transfection (**Figure 2 G, H, H’**, arrows). These are likely a later, secondary consequence of cytokinesis failure, fusion of two or more nuclei (Neumann *et al*., 2010). Lastly, KIF20B knockdown caused a small but significant increase in apoptosis at 24 hours (**Figure 2 I**). Together these data support the conclusion that in dissociated human cells as well as in the developing mouse brain, KIF20B has a role in cytokinesis.

**Figure 2.**
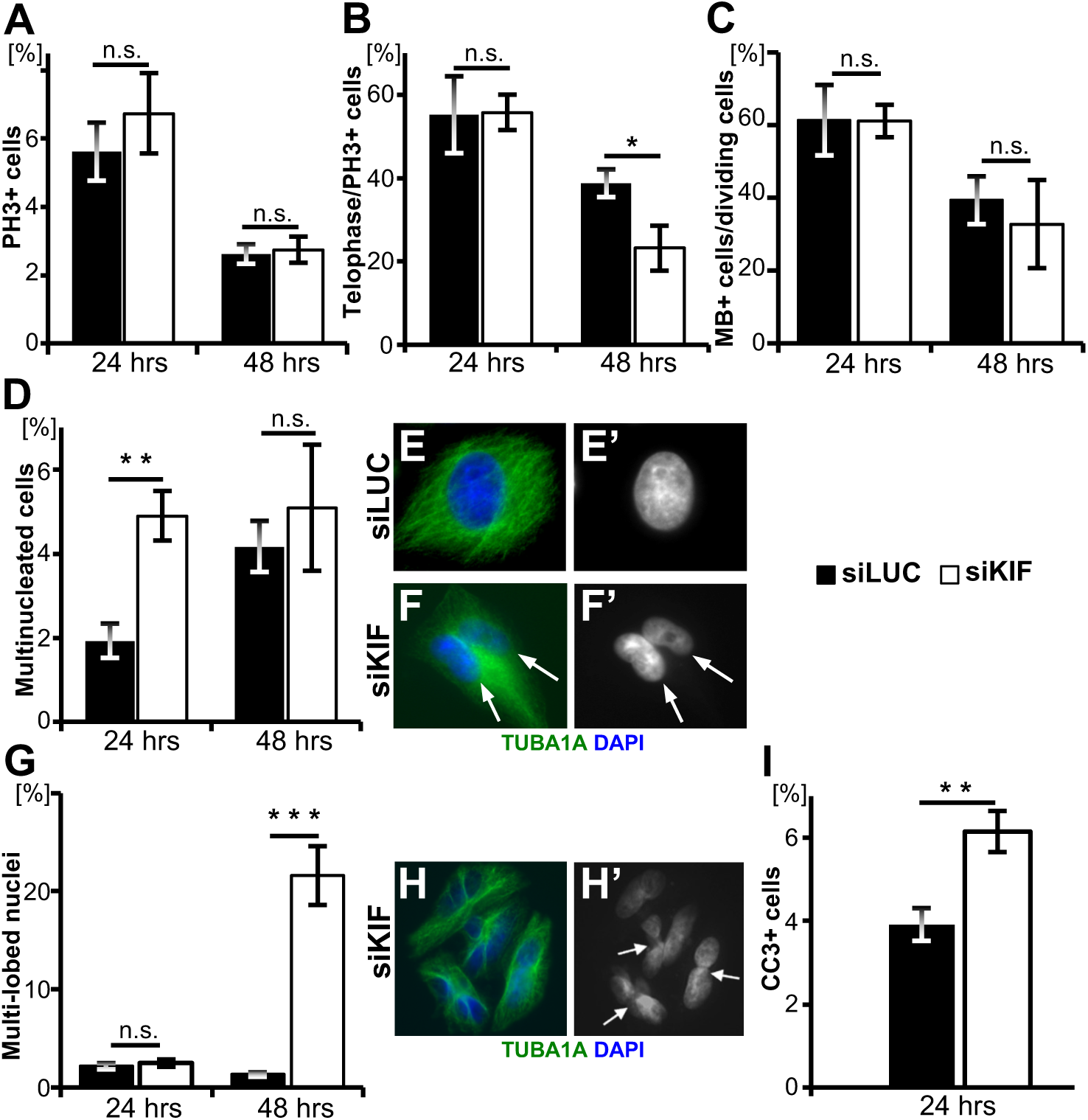
Cytokinesis defects in KIF20B-depleted asynchronous HeLa cell cultures. (A)Average mitotic index of each treatment at 24 hours (n = 5 coverslips/treatment, siLUC = 1192 cells, siKIF = 1408 cells) and 48 hours post-transfection (n = 7 coverslips/treatment, siLUC = 1870 cells, siKIF = 2513 cells). Mitotic index was defined as the sum of all PH3+ cells in Prophase, Metaphase, Anaphase, and Telophase divided by the total cell count. (B)Average percentage of PH3+ HeLa cells in Telophase. Telophase was characterized by the presence of a midbody and condensed chromatin. At 48 hours post-transfection, there was a significantly smaller percentage of Telophase cells (p = 0.0378) in siKIF knockdown. Telophase index was defined as the sum of cells in Telophase out of all mitotic (PH3+) cells at 24 hours (n = 5 coverslips/treatment, siLUC = 67 cells; siKIF = 95 cells) and 48 hours post-transfection (n = 7 coverslips/treatment, siLUC = 49 cells; siKIF = 69 cells). (C)Midbody index was not significantly different in siKIF cells. Midbody index was defined as the sum of cells with midbodies out of the total number of cells quantified at 24 hours (n = 8 coverslips/treatment, siLUC = 2556 cells, siKIF = 3048 cells) and 48 hours posttransfection (n = 6 coverslips/treatment, siLUC = 952 cells; siKIF =1176 cells). (D)At 24 hours post-transfection, there were significantly more multinucleate cells in the knockdown population (p = 0.00468). Quantification of multinucleate cells as a percentage of total cells at 24 hours (n = 5 coverslips/treatment, siLUC = 1192 cells; siKIF = 1408 cells) and 48 hours post-transfection (n = 6 coverslips/treatment, siLUC = 1870 cells; siKIF = 2513 cells). (E and E’) A control siLUC HeLa cell stained with α-TUBULIN (TUBA1A) and DAPI with a single nucleus. (F and F’) siKIF transfected cell stained with TUBA1A and DAPI showing two nuclei within the same cell (white arrows). (G) At 48 hours post-transfection, the percentage of cells with multi-lobed nuclei was significantly greater in the knockdown cells than in the control cells (p = 4.77×10^−5^). Average percentage of cells with multi-lobed nuclei at 24 hours (n = 5 coverslips/treatment, siLUC = 1192 cells; siKIF = 1408 cells) and 48 hours post-transfection (n = 6 coverslips/treatment, siLUC = 1870 cells, siKIF = 2513 cells). (H and H’) siKIF transfected cells stained with TUBA1A and DAPI to show multi-lobed phenotype. White arrows point at constrictions within single nuclei. (I) Average apoptotic index was increased in siKIF cells (p = 0.00639). Average apoptotic index is defined as the percentage of cleaved caspase 3 positive (CC3+) cells at 24 hours post-transfection (n = 8 coverslips/treatment, siLUC = 1878 cells, siKIF = 1486 cells). Coverslips were prepared from 3 independent siRNA transfection experiments.

### Furrow ingression is slower in KIF20B-depleted cells

Next, we sought to analyze cytokinesis in KIF20B-depleted cells in more detail. Analyses were done at 24 hours post-transfection of siRNA, since KIF20B was depleted and phenotypes were already observed. First, we examined the cleavage furrowing stage. In fixed cell images, we noticed that the central spindles of anaphase cells in the siKIF20B-treated cultures sometimes appeared disorganized, with non-parallel microtubule bundles or asymmetric gaps (**Figure 3A, B**, arrowheads). Since the central spindle regulates cleavage furrow positioning and ingression, this notion prompted us to examine the kinetics of furrow ingression in live cell time-lapse imaging. By collecting images every one minute as control or KIF20B-depeleted mitotic cells progressed through cleavage, we observed in each case examined that furrows appeared qualitatively normal, occurring only at the cell equator, ingressing steadily, and never regressing (**Figure 3C and D**). However, there was a quantitative difference. The average total duration of furrow ingression from anaphase onset to completion was significantly increased in KIF20B-depleted cells, from 6.5 minutes to 8.4 minutes (**Figure 3 E**). Plotting furrow width over time demonstrates that the onset of furrow ingression is similar, but ingression proceeds at a slower rate in siKIF20B cells (**Figure 3F**). A smaller but similar reduction was seen in the rate of cell lengthening from pole-to-pole (**Figure 3 G**). Together these data show that while KIF20B is not required for central spindle formation or furrow ingression, it may contribute to central spindle organization and significantly affect the rate of furrow ingression.

**Figure 3.**
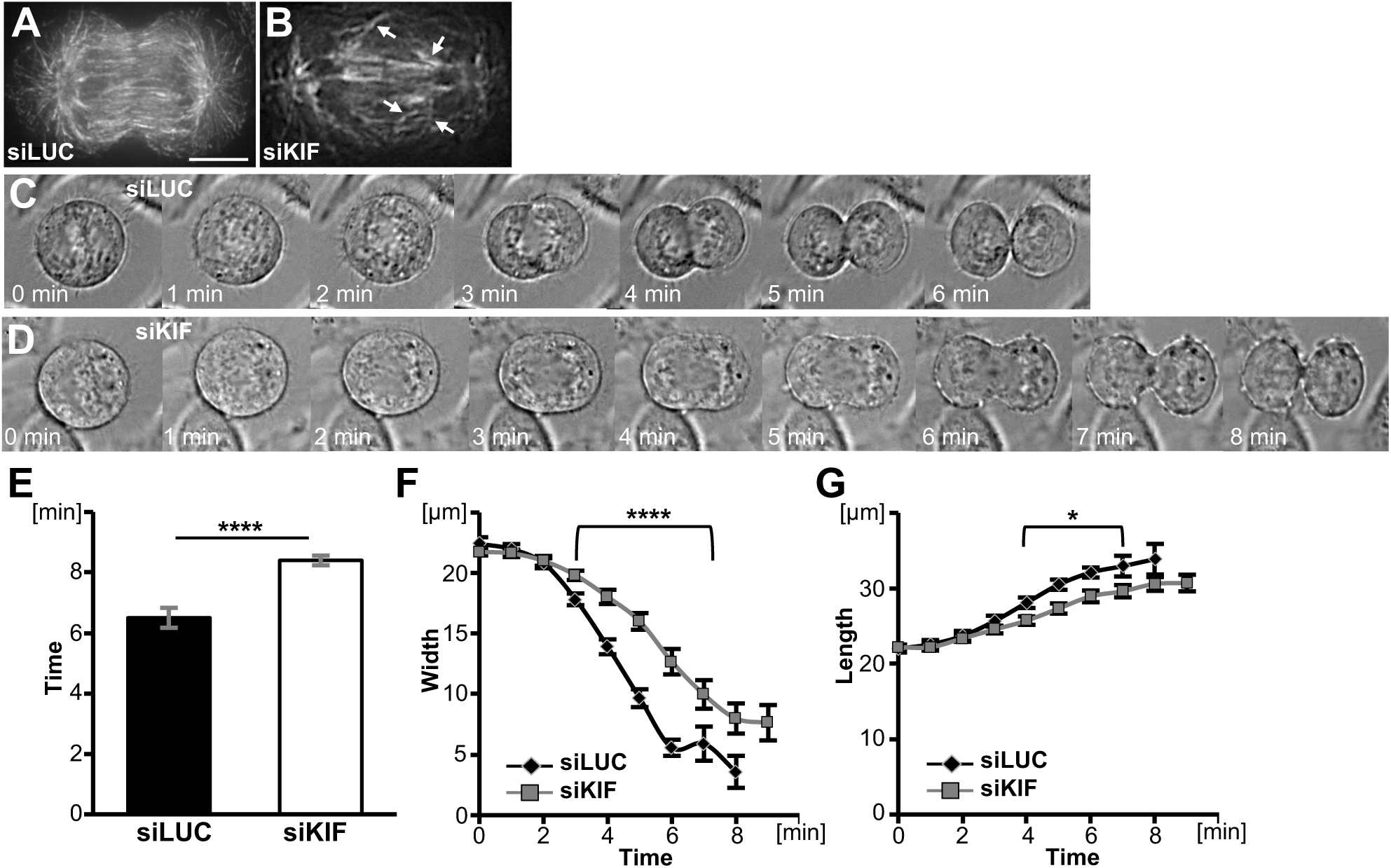
Cleavage furrow ingression is slower in KIF20B-depleted cells. (A,B) Representative images show alpha-tubulin (TUBA1 A) staining of the central spindle in siLUC and siKIF cells in anaphase. Central spindles of siKIF cells have disarrayed microtubules (arrowheads). (C) Representative brightfield time-lapse images of siLUC-treated cell completing furrowing in 6 minutes. (D) Representative siKIF-treated cell completing furrowing in 8 minutes. (E) Average total time from anaphase onset (chromosome segregation onset) to completion of cleavage furrowing ingression was increased in siKIF cells (^∗∗∗∗^p = 3×10^−7^). (F) Furrow diameters of siKIF cells decreased more slowly than those of siLUC cells (for points under bracket, ^∗∗∗∗^p = 3×10^−5^). (G) Cell lengths (pole to pole) of siKIF20B-treated cells increased more slowly than in siLUC cells, (^∗^p ≤ 0.05). n_siLUC_ = 20 cells, n_siKIF_ = 24 cells across 5 independent imaging sessions.

### KIF20B loss alters midbody width but not subdomain structure

We previously showed that in embryonic mouse brains, loss of *Kif20b* disrupted the shapes and positioning of neural stem cell midbodies. Midbodies still formed at the apical membrane of the neuroepithelium, but were more often misaligned, and had an altered distribution of axis ratios, primarily due to increased width (Janisch *et al*., 2013; Janisch and Dwyer, 2016). To test whether this phenotype reflects a cell-autonomous primary requirement for KIF20B in abscission, and whether it occurs in a non-epithelial context, we measured the lengths and widths of midbodies of HeLa cells treated with control or KIF20B siRNA. Indeed, we found that siKIF20B-treated cells did have a significantly shifted distribution of midbody widths, but surprisingly there were more thin midbodies and fewer wide midbodies. This was observed with either tubulin signal or Aurora B kinase immunostaining (**Figure 4 A, B**). Midbodies also tended to be longer when KIF20B was depleted, but the shift was not statistically significant (**Figure 4C**). Together with our previously published midbody analyses in embryonic mouse brains, these data show that loss of KIF20B causes changes in the abscission stage of cytokinesis in a cell autonomous manner.

**Figure 4.**
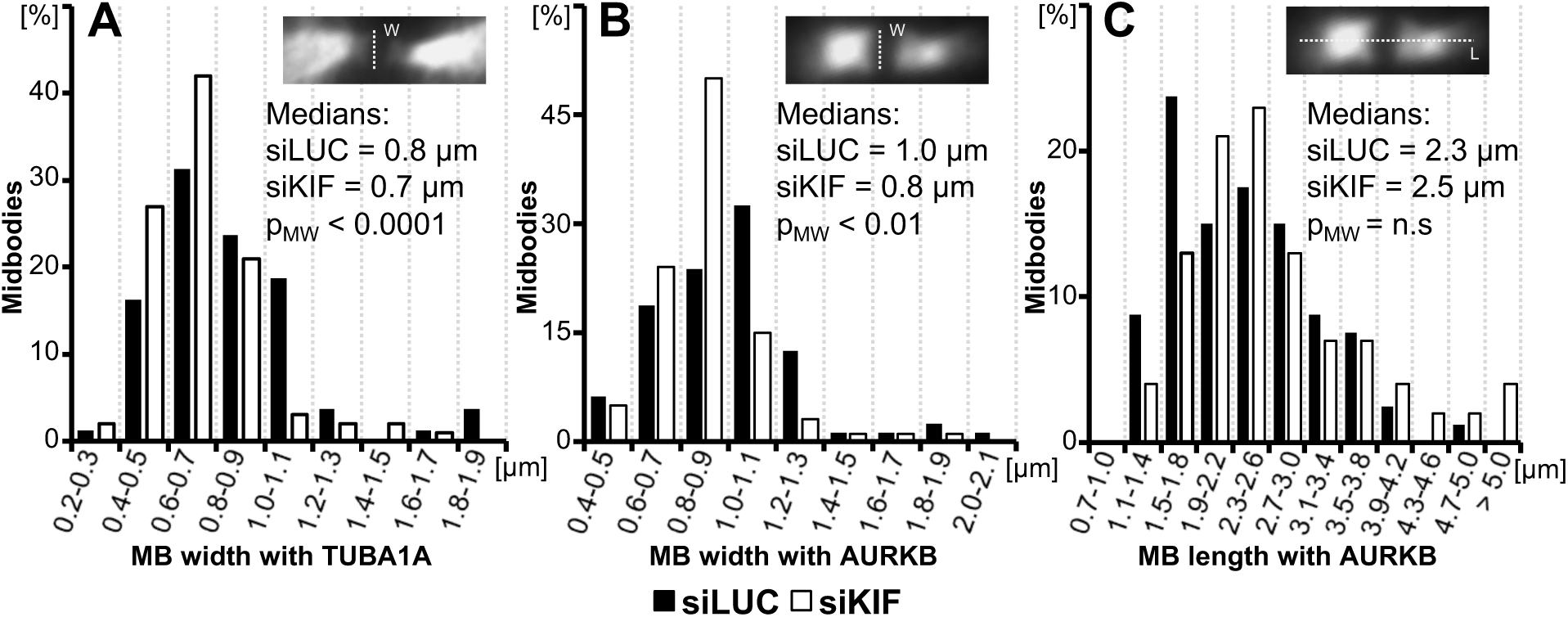
Midbodies tend to be thinner in KIF20B-depleted HeLa cells. 24 hours after transfection with control (siLUC) or KIF20B (siKIF) siRNA, cells were fixed and immunostained for alpha-tubulin and Aurora B kinase (AurkB). Midbodies were imaged and measured for width and length as shown in insets. (A and B) Plots of the distributions of midbody widths for KIF20B-depleted cells (siKIF, white bars) and controls (siLUC, black bars) show that KIF20B-depeleted cells more often have thin midbodies when measured by α-tubulin (TUBA1A) or Aurora B (AURKB) signal (medians: siLUC = 0.8 μm, siKIF = 0.7 μm, p_MW_ < 0.0001, and distribution shape p_KS_ = 0.0205; and siLUC = 1.0 μm, siKIF = 0.8 μm, p_MW_ < 0.01, and distribution shape p_KS_ = 0.000419, respectively). (C) Median midbody length, as measured with Aurora B signal, shows a trend to be increased in siKIF-treated cells, but does not reach statistical significance (p_MW_ = 0.28; p_KS_ = 0.85). p-values (p_MW_) for medians are calculated with Mann-Whitney U-test; p-values (p_KS_) for distribution shape are calculated with Kolmogorov-Smirnov test, n.s. = not significant. n_siLUC_ = 80 midbodies; n_siKF_ = 100 midbodies, from 3 independent siRNA transfections.

We hypothesized that KIF20B may regulate midbody shape directly or indirectly by binding microtubules or by localizing effectors to midbody microtubules. To further investigate midbody structure and protein recruitment when KIF20B is depleted, we took advantage of the wider variety of antibodies that work for immunofluorescence on human cells than on mouse cells. We tested whether several key regulators of cytokinesis are localized properly in the furrow and in subdomains of the midbody (**Figure 5**). Protein Regulator of Cytokinesis 1 (PRC1), which is required for formation of the central spindle (Mollinari *et al*., 2002; Zhu *et al*., 2006) and was shown to interact with KIF20B (Kanehira *et al*., 2007), shows normal localization in KIF20B-depleted cells at the center of the central spindle in anaphase, and in the midbody in two discs in the dark zone as well as the flanks (**Figure 5A-F’**). The Kinesin-6 family member KIF23/MKLP1, required for midbody formation and abscission (Matuliene and Kuriyama, 2002; Zhu *et al*., 2005a) also localizes normally to the central spindle and to the midbody bulge in KIF20B-depleted cells (**Figure 5 G-L”**). Aurora B kinase (AURKB) localizes to the central spindle and the flanks of the midbody in both control and siKIF20B-treated cells (**Figure 5 G-L”**). Activated Aurora B kinase phosphorylated at T232 (pAURKB), which was shown to regulate the abscission checkpoint (Steigemann *et al*., 2009; Carlton *et al*., 2012), localizes to the center of the midbody in both control and KIF20B-depleted cells (**Figure 5 M-R’**). Anillin (ANLN), a scaffold for recruiting furrow proteins, appears normally localized to the furrowing membrane, as well as in midbodies as a ring around the center at early stages, and at constriction sites at late stages (**Figure 5 S-X’)**. Alpha-actinin-4 (ACTN4), which regulates furrowing (Mukhina *et al*., 2007), was pulled down as a candidate binding protein with KIF20B (Maliga *et al*., 2013). However, it appears indistinguishable in the furrows of KIF20B-depleted cells, and is not enriched in midbodies in control or KIF20B-depleted cells (**Figure 5 Y-DD’)**. These data demonstrate that when KIF20B is depleted, several key regulators of furrowing and abscission are localized normally, and that the primary midbody subdomains of dark zone, bulge, and flanks are specified.

**Figure 5.**
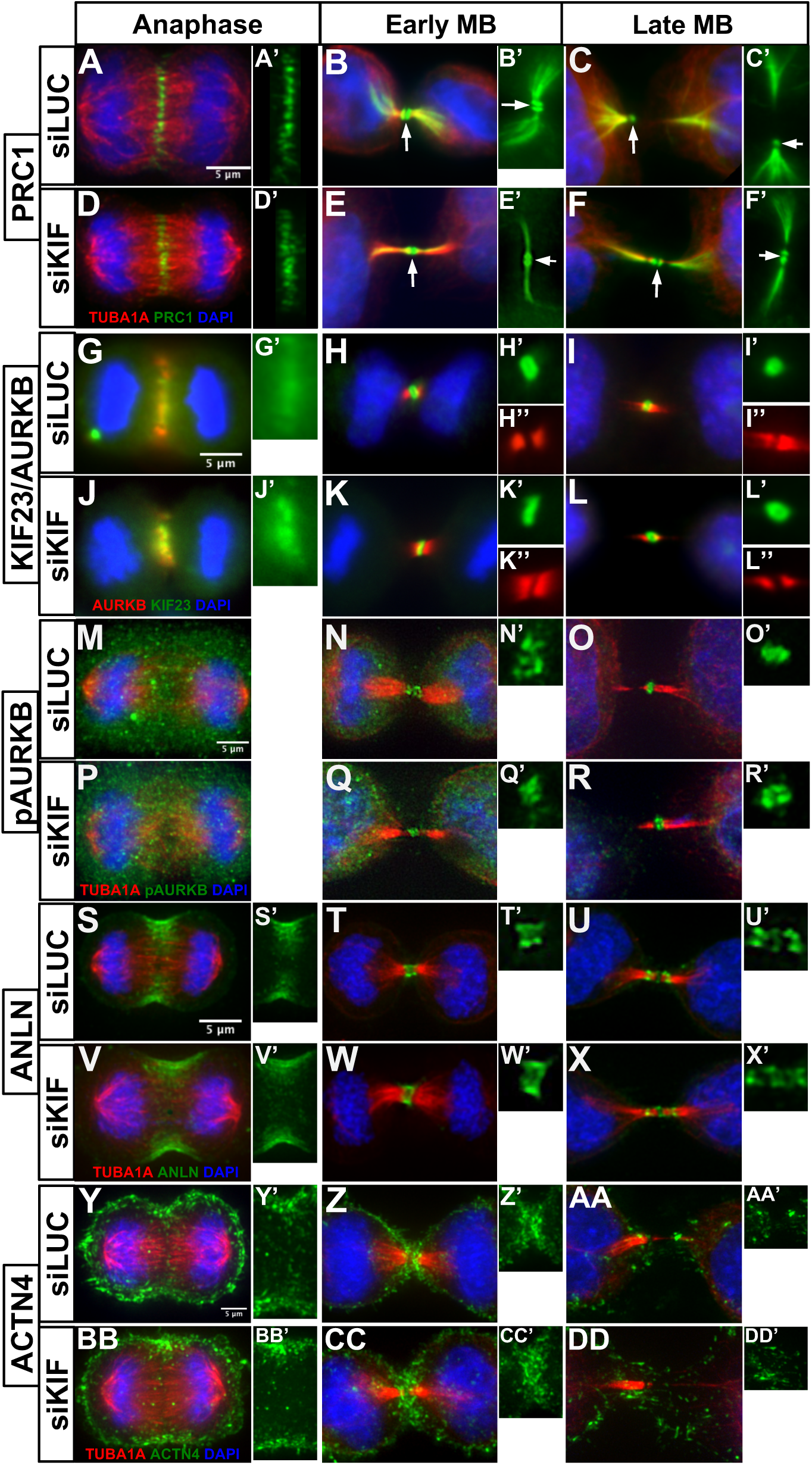
Midbody assembly appears to occur normally in KIF20B-depleted cells. (A, A’ and D, D’) PRC1 localizes to the central spindle in anaphase in both siLUC and siKIF cells. (B, B’ and E, E’) In early midbody (MB) stage, PRC1 lines the microtubules (MT) of the midbody flanks and reaches into the cell on the microtubule network. It also forms two distinct discs around the dark zone of the midbody (white arrow). (C, C’ and F, F’) In late midbody stage, PRC1 can still be found on the midbody flank microtubules reaching into the cell. (G, G’ and J, J’) MKLP1 localizes to the central spindle in anaphase in siLUC and siKIF cells. It can be found in the dark zone of the midbody both in early (H, H’ and K, K’) and late midbody stage (I, I’ and L, L’). (H’’, I’’ and K”, L”) Aurora Kinase B (AURKB) localizes to the flanks of the midbody in early and late midbody stages in siLUC and siKIF cells. (M and P) pAURKB is diffusely localized during anaphase in siLUC and siKIF cells. In early (N and Q) and late (O and R) midbody stage, pAURKB localizes to the dark zone of the midbody. In the early stage (N’ and Q’) it forms a diffuse blob, whereas it appears to be disc-shaped in the later stages (O’ and R’) in siLUC and siKIF cells. (S, S’ and V, V’) Anillin (ANLN) localizes to the furrow during anaphase in siLUC and siKIF cells. In early midbody stage (T and W), ANLN forms a sheet-like ring (T’ and W’) around the dark zone of the midbody. In late midbody stage (U and X), ANLN can be found in the dark zone of the midbody as well as at the constriction sites (U’ and X’). (Y, Y’ and BB, BB’) α-actinin-4 (ACTN4) encases the two daughter cells but clearly accumulates in the cleavage furrow during anaphase in both siLUC and siKIF cells. (Z, Z’ and CC, CC’) In early midbody stage, ACTN4 enriches in a half-circle shape at the edges of both daughter cells underneath the midbody. (AA, AA’ and DD, DD’) In late midbody stage, the enrichment of ACTN4 around the midbody disperses in both siLUC and siKIF cells. Images taken with a widefield microscope (G - L) or DeltaVision deconvolution microscope (A - F’; M - X’). At least 15 midbody-stage and 5 anaphase cells imaged for each marker and condition. Scale bars are 5 μm for all images.

### KIF20B-depleted midbodies have reduced microtubule constriction sites and increased CEP55 intensity

Since the higher proportion of thin midbodies in KIF20B-depleted cells suggested a problem in abscission (**Figure 4**), but the major midbody subdomains appear to assemble normally (**Figure 5**), we sought to further characterize the late stage midbodies. First, to address whether midbodies mature properly, we quantified microtubule constriction sites. In fixed midbodies immunostained for tubulin, constriction sites appear as pinches or gaps in microtubule signal, and are the presumed abscission sites (Guizetti *et al*., 2011; Hu *et al*., 2012). These constriction sites (also called secondary ingressions) may be visible on one or both sides of the midbody bulge/dark zone (**Figure 6 A, B**). Interestingly, we found that in the KIF20B-depleted cells, a smaller percentage of midbodies had constriction sites (**Figure 6 C**). This suggests a defect in formation or structure of constrictions, an important aspect of midbody maturation in preparation for abscission. As a second approach to assess midbody maturation, we analyzed the localization of CEP55, which is a key abscission regulator that accumulates in late midbodies, starting about 50-60 minutes after anaphase onset in HeLa cells. It localizes to two rings within the dark zone and then recruits ESCRT proteins to mediate abscission (Bastos and Barr, 2010). Similarly, we observed that in midbodies without constriction sites, Cep55 immunostaining signal appeared in two discs perpendicular to the microtubules inside the dark zone, resolvable with or without deconvolution (**Figure 6 D, F, H, J**). In late midbodies with at least one microtubule constriction site, the CEP55 discs appeared closer together, usually still resolvable with deconvolution in most control cells, but less often in KIF20B-depleted cells (**Figure 6 E, G, I, K**; arrowheads point to constriction sites). Interestingly, in KIF20B-depleted midbodies with at least one constriction, the CEP55 signal had a greater maximum intensity, though occupying the same area (**Figure 6 L, M**). These findings of reduced constriction sites and increased CEP55 intensity suggest that KIF20B-depleted midbodies may have defects in late steps of maturation, or in temporal regulation of the events leading to abscission.

**Figure 6.**
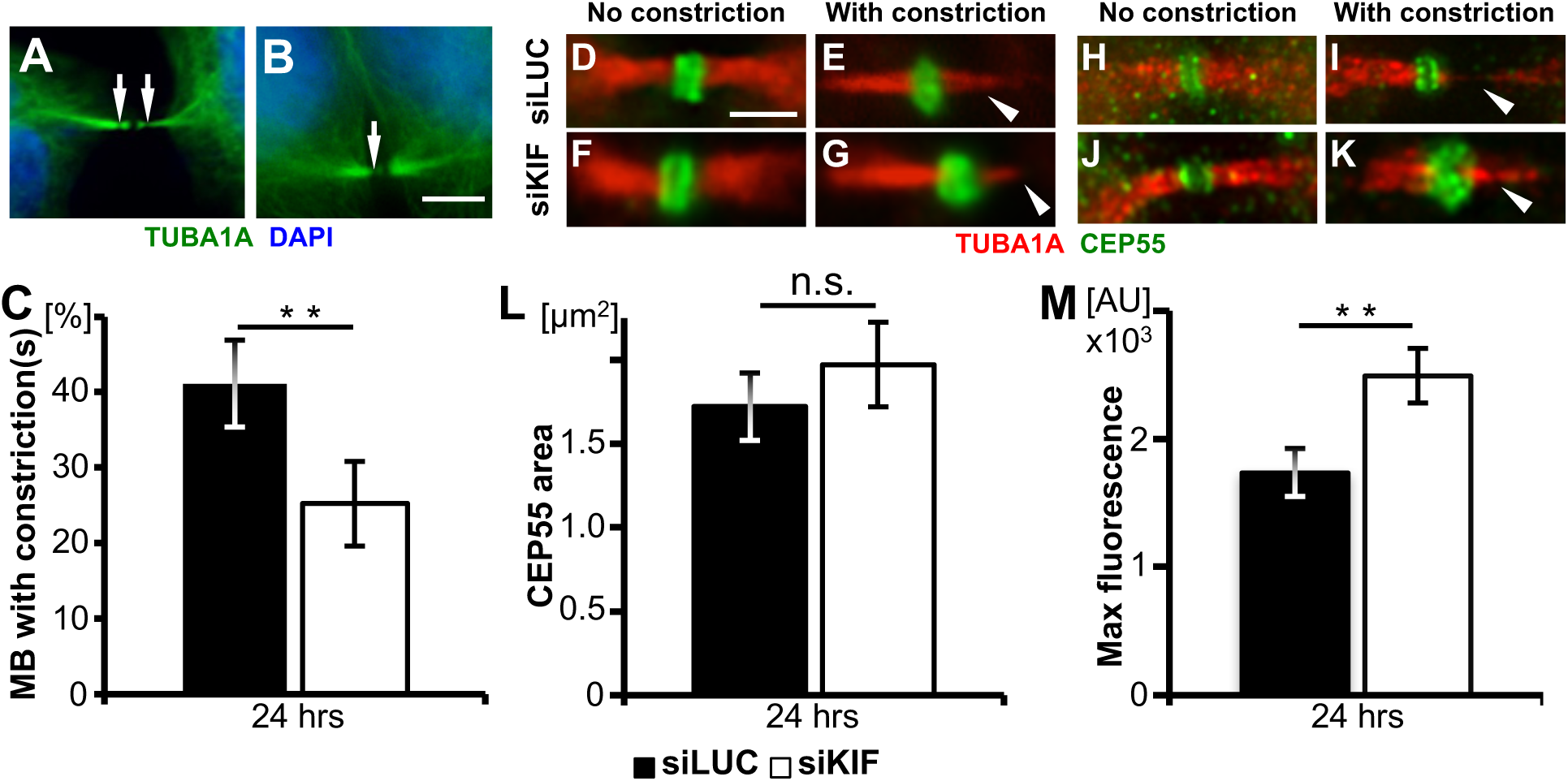
Midbodies of KIF20B-depleted cells show defects in late stages of maturation. (A and B) Representative images of tubulin in midbodies with one or two constriction sites visible (arrows). (C) The average percentage of midbodies having at least one constriction site was reduced in KIF20B-depleted cells at 24 hours post-transfection (p = 0.013). (D - K) CEP55 localizes to the dark zone of the midbody in both siLUC and siKIF20B-treated cells. D - G show non-deconvolved images and H - K show deconvolved images. In midbodies without constriction sites, CEP55 appears in two distinct discs. In late midbodies having constriction sites (arrowheads in E, G, I, K), CEP55 is more dense but can still be resolved into two discs by deconvolution is control cells (I) but not in siKIF-treated cells (K). (L) The average area of CEP55 signal in the dark zone of late midbodies with constriction sites is not significantly different between siLUC and siKIF-treated cells. (M) The average maximum fluorescence intensity of CEP55 is significantly higher in late midbodies with constriction sites after KIF20B depletion. All p-values were calculated with two-tailed Student’s t-test, except in (C) paired t-test was used, n = 5 coverslips/ treatment from 3 independent transfections. Scale bars for A and B, 5 μm; for D – K, 2.5 μm.

### KIF20B depletion disrupts abscission timing

To test directly whether loss of KIF20B causes delays or failures in abscission, we performed live cell time-lapse imaging. First, the cell-permeable tubulin dye Silicon-Rhodamine Tubulin (SiR-tubulin) was used to monitor abscission (**Figure 7A, B**). This far red dye only fluoresces when bound to polymerized microtubules (Lukinavicius *et al*., 2014). Figures 7A and B show examples as imaged by wide-field microscopy at lower (**7A**) or higher magnification (**7B**). In both cases, the first abscission (a1, white arrowhead) and second abscission (a2, yellow arrowhead) can be observed, and the midbody remnant can be followed for at least an hour after abscission (**7B**, open arrow at 105 minutes). Interestingly, at the higher magnification, the severed midbody flanks can also still be resolved as coherent microtubule bundles more than an hour after the second abscission (**Figure 7B**, wide arrows at 105 minute time point.).

**Figure 7.**
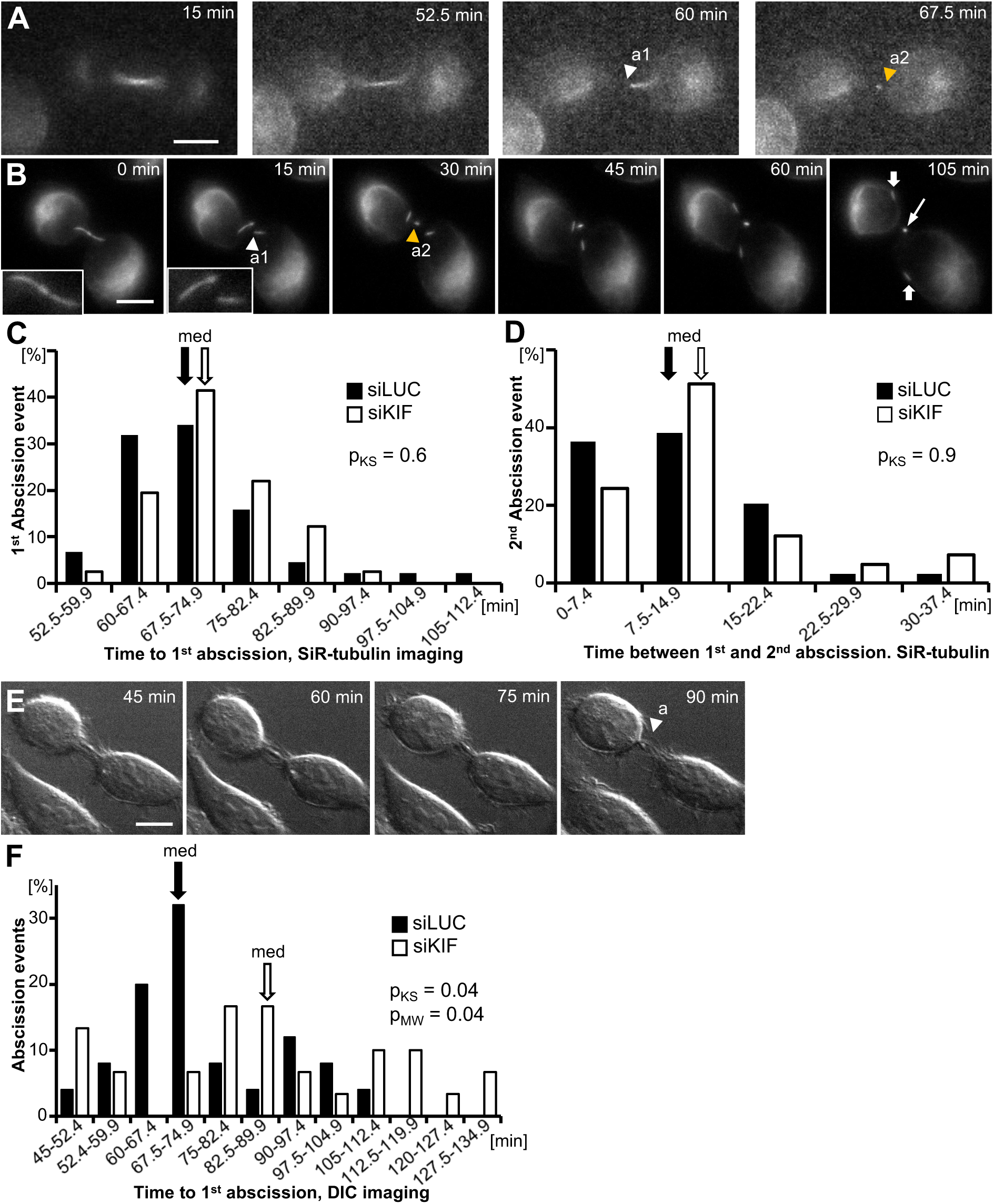
KIF20B depletion delays abscission. (A and B) Example widefield time-lapse imaging of abscission in a HeLa cell labeled with SiR-Tubulin at 20x (A) and 63x (B) (arrowhead a1: abscission 1, arrowhead a2: abscission 2, filled arrow: midbody flanks, open arrow: midbody remnant). Images captured every 7.5 minutes (A) and every 15 minutes (B). Insets in B are showing the intact midbody and the first abscission. (C) Frequency distribution of time from anaphase to first abscission shows a similar median time (67.5 minutes, arrows, p = 0.13 (M-W)) and similar distribution shape (K-S test p=0.6) for siLUC (Black bars) and siKIF-treated cells (white bars) imaged with SiR-tubulin. n = 44 siLUC cells and 41 siKIF20B cells from 4 experiments. (D) The median time between first and second abscissions (arrows, 7.5 minutes, p_M-W_= 0.09) imaged with SiR-tubulin was not different in KIF20B-depleted cells (siKIF) and controls (siLUC). The distributions were not significantly different between siLUC and sKIF-treated cells (p_K-S_=0.9). Sample is same as in C. (E) Representative selected planes from time-lapse z-stack images of abscission using differential interference contrast (DIC) microscopy, captured every 7.5 minutes (arrowhead a signifies abscission). The second abscission event was not discernible by DIC imaging. (F) Frequency graph of time from anaphase to first abscission discerned using DIC time-lapse microscopy shows both an altered distribution of times and an increased median time (black arrow versus white arrow) in siKIF-compared to siLUC-treated cells. (distribution p_K-S_ =0.04; medians 67.5 minutes versus 82.5 minutes, p_M-W_=0.04). n = 25 siLUC cells and 30 siKIF cells from 4 experiments. K-S: Kolmogorov–Smirnov test, M-W: Mann-Whitney U-test. Scale bars, 10 *μ*m.

By imaging every 7.5 minutes, we found that control cells had a median time from anaphase onset to first abscission of 67.5 minutes (**Figure 7C**, black bars). This is similar to the timing observed previously in HeLa cells using both GFP-tubulin or photoactivatable dye transfer (Steigemann *et al*., 2009; Guizetti *et al*., 2011), confirming that SiR-tubulin is a valid tool for abscission studies. Surprisingly, however, in KIF20B-depleted cells, the median time to first abscission was not significantly different from controls (**Figure 7C**, white bars). Since midbody inheritance or release has been proposed to influence daughter cell fate in stem cell divisions (Dubreuil *et al*., 2007; Ettinger *et al*., 2011; Kuo *et al*., 2011; Salzmann *et al*., 2014), we also wanted to test whether KIF20B regulates the time between first and second abscissions. We found that the second abscission usually occurred within 15 minutes of the first abscission (~75% of the cells), in both siLUC and siKIF-treated cells (**Figure 7D**). This is similar to but slightly faster than what was reported using GFP-tubulin (Guizetti *et al*., 2011; Gershony *et al*., 2017). Interestingly, the cells that had the longest times to first abscission were not the ones that had the longest times between first and second abscissions, suggesting they are independent events (data not shown).

The lack of a detectable change in abscission timing upon KIF20B depletion was surprising, given the late midbody defects observed. Since SiR-tubulin is a derivative of docetaxel, which stabilizes microtubules, we hypothesized that SiR-tubulin might be rescuing the effect of loss of KIF20B by stabilizing microtubules. To test this hypothesis, we employed a different method of scoring abscission, differential interference contrast (DIC) imaging, without SiR-tubulin (**Figure 7E**). With this method, only the first abscission event could be discerned with confidence. The median time to first abscission in control siRNA cells was 67.5 minutes, the same as seen with SiR-tubulin (**Figure 7C and 7F**, black arrows). Remarkably, however, we found that in siKIF20B-treated cells, abscission timing was significantly dysregulated, with a wider distribution of times, and a median increase of 15 minutes (**Figure 7F**, white bars, white arrow). Together these live cell experiments show that KIF20B regulates the timing of abscission, and further suggest that KIF20B does so by stabilizing microtubules.

## DISCUSSION

With the major players required for cytokinesis largely identified, there is a growing need to understand the roles of proteins that regulate the temporal aspects of cytokinesis, or play specialized roles in different cell types. We have demonstrated here that KIF20B, a Kinesin-6 family member required for normal brain size, temporally regulates both cytokinetic furrow ingression and abscission. Furrow ingression was slower, and abscission time was dysregulated and increased. KIF20B does not appear to be required for midbody assembly or its subdomain structure, but may be involved in late steps of maturation such as creating microtubule constriction sites.

Multiple pieces of data suggest that one function of KIF20B is to crosslink or stabilize tight microtubule bundles. Other groups showed that KIF20B is sufficient to crosslink microtubules in a cell-free assay (Abaza *et al*., 2003), and that pavarotti/MKLP1 inhibits microtubule sliding in *Drosophila* neurons (del Castillo *et al*., 2015). We previously showed that in growing axons of *Kif20b* mutant neurons, microtubules are less tightly packed and frequently invade growth cone filopodia (McNeely *et al*., 2017). Here and in prior work we showed that midbody width distributions are altered when KIF20B is absent, and microtubule constriction sites are less detectable. In addition, the finding that SiR-tubulin appears to rescue the abscission timing defect also supports the idea that KIF20B may help stabilize microtubules in preparation for abscission.

The phenotypes caused by depletion of KIF20B from HeLa cells are subtle compared to those seen from depletion of other midbody proteins or the other Kinesin-6 family members. By contrast, HeLa cells depleted of KIF23/MKLP1 usually fail to assemble a proper midbody core, regress their furrow, and become binucleate (Matuliene and Kuriyama, 2002; Zhu *et al*., 2005b). HeLa cells depleted of KIF20A/MKLP2 may either fail to complete furrow ingression (Neef *et al*., 2003; Kitagawa *et al*., 2013), or assemble a midbody but fail to complete abscission (Zhu *et al*., 2005b). This may explain why KIF20B was not identified in high throughput RNAi screens for genes involved in cell division (Kittler *et al*., 2004; Kittler *et al*., 2007; Neumann *et al*., 2010), or for motor protein knockdown phenotypes (Zhu *et al*., 2005b). In retrospect, the magnitude of KIF20B phenotypes probably did not meet the thresholds applied in these screens, but are only revealed upon focused quantitative analysis. Our developmental screen, rather than a single-cell screen, revealed the importance of this Kinesin-6 (Dwyer *et al*., 2011).

Although KIF20B is not absolutely required for cytokinesis, our previous analyses showed that it is essential for normal brain size in the mouse. The mutant embryos grow and form a brain, suggesting that at least some tissues can compensate for the loss of *Kif20b*, and that many neural stem cell divisions occur normally. However, the neural stem cell midbodies in the cerebral cortex show abnormal shapes and organization, and there is increased apoptosis (Janisch *et al*., 2013). What we do not yet know is whether the changes in cytokinesis observed upon KIF20B depletion in HeLa cells are increased in magnitude in neuroepithelial stem cells, or whether the neural cells are simply much more sensitive to even slight inefficiencies in cytokinesis. Polarized neuroepithelial stem cells are very tall and thin, and furrow ingression proceeds basal to apical. In addition, furrowing and abscission must be coordinated with the inheritance of apical adhesions and cell fate determinants, and delamination of neuronal daughters. These factors suggest that cytokinesis in a polarized epithelium is more challenging than in a HeLa cell and requires both spatial and temporal precision (Dwyer *et al*., 2016; Johnson *et al*., 2017). Interestingly, in the neuroepithelium, Kif20b is important for keeping apical midbodies aligned with apical adherens junctions (Janisch *et al*., 2013). A slight delay in abscission might trigger apoptosis; or dysregulation of abscission timing may disrupt the control of symmetric versus asymmetric divisions. Even a small percent of stem cell death can have a large effect due to the loss of a lineage of progeny over developmental time. Thus, HeLa cells, the most commonly used mammalian system to study cytokinesis, cannot fully model the cytokinesis mechanisms and phenotypes of developing epithelia like the early brain.

Despite such limitations, the cell line work herein has generated hypotheses about KIF20B’s roles in cytokinesis to test in the developing brain. Future experiments will examine whether similar defects in midbody maturation and abscission occur in neural stem cells in culture and in intact brain explants, as well as how the timing of furrowing and abscission are coordinated with tissue polarity and daughter fates. This work underscores the need for more studies of cytokinesis in developing tissues in addition to isolated single cells. There may be many important players that regulate the timing or precision of cytokinesis that have not yet been identified from single cell screens. Mutations in these genes may be discovered as causing developmental or functional defects in animal models or human clinical diseases.

## MATERIALS AND METHODS

### Cell Culture

HeLa cells (human cervical carcinoma cells) were obtained from ATCC; we did not re-authenticate. HeLa cells were grown in 10 cm petri dishes with DMEM (Gibco, ThermoFisher Scientific, MA, USA) supplemented with 10% FBS (Atlanta Biologicals, GA, USA), 15 mM HEPES (Gibco, ThermoFisher Scientific, MA, USA), 1X glutamine (Gibco, ThermoFisher Scientific, MA, USA), 1X non-essential amino acids (Gibco, ThermoFisher Scientific, MA, USA) and 1X Penicillin/Streptomycin (Gibco, ThermoFisher Scientific, MA, USA). Cells were grown at 37 °C and 5% CO_2_ until they reached ~70% confluency before subculturing for usage in experiments. Coverslips were examined for mycoplasma contamination monthly by DAPI staining of nuclei. For siRNA transfections, unsynchronized HeLa cells were harvested, counted and plated onto 22 mm diameter glass coverslips with a density of 50000 cells per coverslip, unless otherwise stated.

### Transfection with GFP-KIF20B

GFP-KIF20B plasmid was provided by F. Pirollet (Abaza *et al*., 2003), and verified by sequencing. Cells were used when ~70% confluent and transfected using PolyJet transfection kit (SignaGen Laboratories^®^, MD, USA) according to the manufacturer’s instructions. Cells were co-transfected with mCherry to allow for transfection efficiency. After 24 hrs, cells were fixed with 4% PFA followed by ice-cold methanol prior to staining and imaging.

### Transfection with siRNA

For knockdown experiments, we independently tested two custom-made small interfering RNAs (Invitrogen) previously reported to knock down human KIF20B (siRNA#1: 5’ AAAGGACAGAGUCGUCUGAUUUU, siRNA#2: 5’ AAUGGCAGUGAAACACCCUGGUU from (Abaza *et al*., 2003)). Since we observed that siRNA#2 depleted KIF23/MKLP1 as well as KIF20B, we discontinued use of siRNA#2 and used siRNA#1 for all experiments shown. A standard negative control siRNA to firefly luciferase (siLUC) was designed by Invitrogen. Cells were transfected using Lipofectamine^®^ RNAiMAX (Thermo Fisher) transfection reagent according to the manufacturer’s instructions. The final concentrations of siRNA were 10 nM. Cells were either fixed 24 hours or 48 hours (with medium change) after transfection and analyzed by immunofluorescence staining.

For live cell imaging, cells were plated at a density of 50,000 cells per chamber in a two-chamber coverglass (Nunc Lab-Tek, ThermoFisher Scientific, MA, USA) before transfection and imaged 22 – 24 hrs after transfection.

### Immunocytochemistry

The standard fixation for most antibodies used was 4% PFA/PBS for 2 min at room temperature, followed with −20 °C methanol for 10 min. Cells were then washed 3X with PBS to remove any residual methanol and stored at 4 °C until usage. For phospho-AURORA kinase B (pAURK) staining, cells were fixed with −20 °C methanol for 10 min followed by 3 washes with PBS. For staining for ANILLIN (ANLN) and alpha-ACTININ 4 (ACTN4), cells were fixed with 10% TCA in cytoskeleton buffer with sucrose (CBS; 10 mM MES pH 6.1, 138 mM KCl, 3 mM MgCl_2_, 2 mM EGTA, 0.32 M sucrose). Shortly, 10% TCA in CBS was added to the cells and incubated on ice for 15 min. Cells were permeabilized with 0.2% Triton-X100, 50 mM glycine in PBS for 2 min on ice, then quenched with 50 mM glycine in PBS for 20 min.

Cells were blocked with 2% normal goat serum (NGS) in PBS with 0.1% Triton-X 100 (PBST) for 1 hour at room temperature. After 3 washes with PBS for 10 min each, primary antibodies diluted in blocking buffer were applied for 3 hours at room temperature. Appropriate secondary antibodies diluted in blocking buffer were applied after 3 washes with PBS for 5 min each and incubated for 30 min at room temperature in the dark. Cells were mounted with flouromount (Diagnostic BioSystems, CA, USA) after a nuclear counterstain with DAPI (Fisher Scientific, PA, USA) and 2 washes with PBS for 10 min.

### Antibodies

Primary antibodies used were as follows: mouse monoclonal DM1alpha (alpha-tubulin) (1:500) was from Abcam (MA, USA); rat anti-TUBA1A (clone YL ½) (1:750) was from Novus Biologicals (CO, USA); mouse polyclonal anti-CEP55 (1:200) was from Abnova (CA, USA); mouse anti-Aurora kinase B (AURKB) (1:300) was from BD Biosciences (MA, USA); rabbit anti-phospho-Aurora kinase B (pAURKB) (1:200) was from Rockland (PA, USA); goat anti-anillin (ANLN) (1:300); rabbit anti-PRC1 (1:50) and mouse anti-MPP1(KIF20B) (1:300) were from Santa Cruz (CA, USA); rabbit anti-mouse-Kif20b (1:500) was custom-made by Covance (NJ, USA; (Janisch *et al*., 2013)); rabbit anti-cleaved caspase 3 (CC3) (1:200) and rabbit anti-phosphohistone H3 (PH3, Alexa Fluor 647 conjugated) (1:400) were from Cell Signaling (MA, USA); rabbit anti-alpha-ACTININ4 (ACTN4) (1:250) was from Millipore (MA, USA). The monoclonal mouse anti-MPP1(KIF20B) antibody was validated by verifying that the midbody staining was lost in *Kif20b* mutant mouse cells. All other primary antibodies were validated by verifying that the staining patterns matched multiple published reports. Secondary antibodies were goat or donkey polyclonal IgG (H+L) conjugated to Alexa fluorophores against according species and were used at 1:200 (Life technologies, NY, USA).

### Imaging and data analysis

**Fixed images** were either collected with a Zeiss AxioVision ImagerZ1 widefield microscope with 40x/1.3 or 100x/1.25 Oil M27 APO objectives or a DeltaVision Elite with TrueLight deconvolution microscope with 60x/1.42 Oil Plan APO objective. For comparisons between siLUC and siKIF20B knockdown cells, exposure times were kept constant for each treatment and images were taken on the same day. For DeltaVision images, deconvolved maximum intensity projections of z-stacks are displayed unless otherwise specified. For image analysis and data acquisition of images, we used Fiji/ImageJ (http://imagej.net/Fiji/Downloads).

**Live cell imaging** was done on an inverted Zeiss AxioObserver microscope equipped with a temperature and CO_2_ -controlled chamber set at 5% CO_2_ and 37°C, multipoint acquisition and DefiniteFocus. Illumination and exposure times were kept to minimum practical levels. Images were acquired with an AxioCam Mrm camera and Zeiss Zen software. Cells in metaphase were chosen in brightfield based on their shape and chromatin appearance. siKIF20B cells were imaged first, followed by siLUC cells. Time 0 was the last frame when chromatin was aligned at the metaphase plate. Cells were unsynchronized.

**Live imaging of furrow ingression** was done by collecting brightfield images with a 63x oil objective. Image stacks with 0.75 um step size were captured every minute. Furrow was considered completely ingressed when no further ingression could be observed using raw z-stacks or projections. In no case was furrow regression observed.

#### For abscission data

##### Brightfield

Image stacks with 0.5um step size were collected every 7.5 min with a 63x oil objective and differential interference contrast (DIC) for 4-6 hours. Abscission was scored in the z-plane where the midbody bulge was visible, and was defined as the separation of the midbody from one or both cells (i.e. movement of the bulge away from a cell, and connection no longer detectable). 25 siLUC cells and 30 siKIF20B cells were analyzed.

##### SiR-tubulin fluorescent imaging

Cells were treated with 100 nm Silicon-Rhodamine Tubulin (SiR-Tubulin) 6 hours after transfection according to the manufacturer’s instruction (cytoskeleton.com, CO, USA) and at least 6 hours before imaging. Cells were confirmed to be in metaphase and to have bipolar spindles using the SiR-Tubulin. Cells with more than two spindles (observed in both control and depleted cultures) were not used for measurements. Image stacks with 1.0 um step size were captured with a 20x objective and a Cy5 (far red) filter, and maximum intensity projections of selected planes were used for analysis. The first abscission was defined as the first microtubule break, with no visible fluorescence between the flank and the bulge. The second abscission was defined as the last break of the microtubules allowing the release of the midbody remnant into the medium. SiR-Tubulin enabled the visualization of both abscission events. DIC images were blinded for analysis.

Live cell images were collected and analyzed using Zen Blue software (Zeiss).

**Other data analysis and statistics** were done with Microsoft Excel, GraphPad Prism, and PAST. (http://palaeo-electronica.org/2001_1/past/issue1_01.htm). Unless otherwise indicated, p-values were calculated with two-tailed Student’s t-test, and error bars are S.E.M.

## ACKNOWLEDGEMENTS

We thank Fabienne Pirollet for the GFP-KIF20B plasmid. We are grateful to Todd Stukenberg, Jim Casanova, Bettina Winckler, Jing Yu, Xiaowei Lu, and members of their labs for advice and discussions. This work was supported by the National Institutes of Health (R01 NS076640 to NDD), an American Cancer Society seed grant (ACS-IRG-81-001-26 to NDD), and a Harrison Undergraduate Research Award to JMD.

---

**Supplemental Figure 1.**
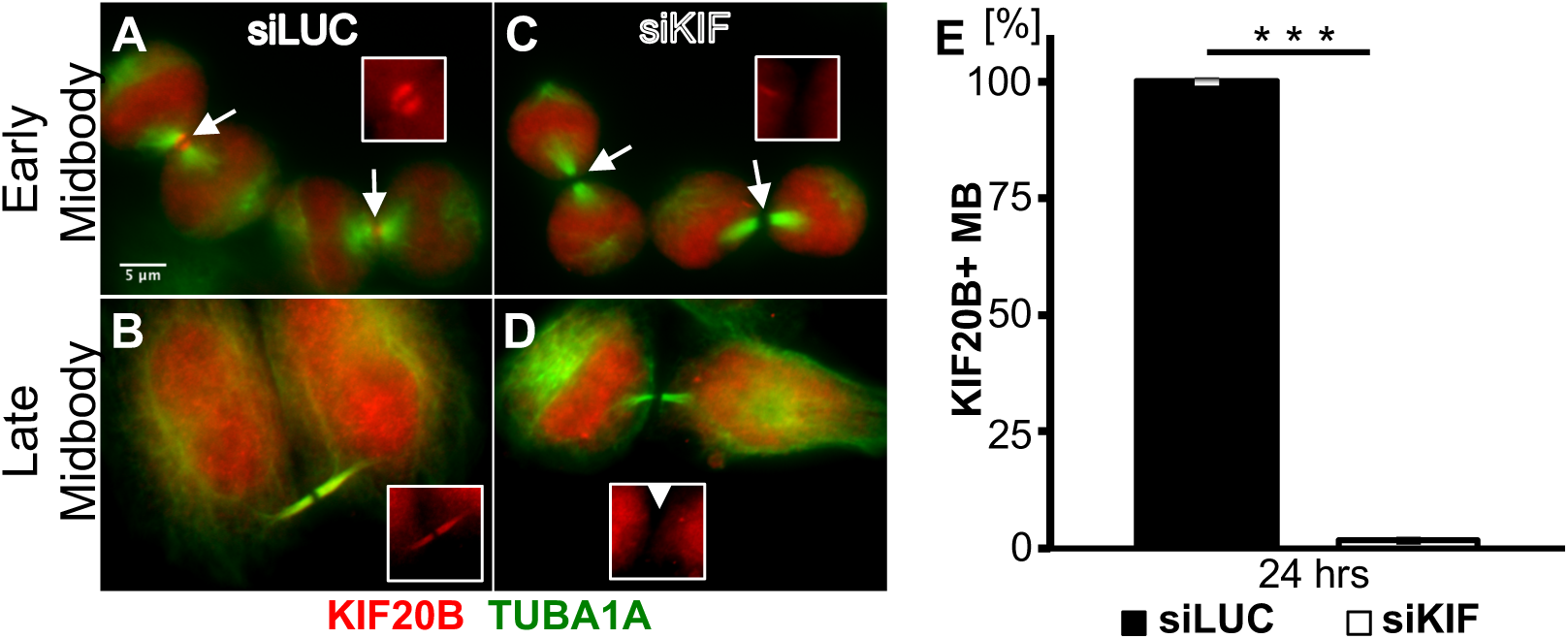
Knockdown efficiency of siRNA to human KIF20B. (A and B) In cells treated with control siRNA to the Luciferase gene (siLUC), endogenous KIF20B signal flanks the dark zone of early midbodies (A, white arrows) and is more spread out in the midbody at later stage (B, white arrowheads). (C and D) In cells treated with siRNA to KIF20B (Abaza 03), KIF20B is not detected in the midbodies at early (white arrows) or late (white arrow heads) stages. Insets show KIF20B staining (red). (E) Quantification of siKIF20B RNA knockdown efficiency results in a nearly 100% depletion of detectable KIF20B in the midbody. p < 0.001, n = 3 coverslips/treatment, 71 midbodies for siLUC and 60 midbodies for siKIF. Scale bar is 5 *μ*m.

